# SpaceBar enables clone tracing in spatial transcriptomic data

**DOI:** 10.1101/2025.02.10.637514

**Authors:** Grant Kinsler, Caitlin Fagan, Haiyin Li, Jessica Kaster, Maggie Dunne, Robert J. Vander Velde, Ryan H. Boe, Sydney Shaffer, Meenhard Herlyn, Arjun Raj, Yael Heyman

**Affiliations:** Department of Bioengineering, School of Engineering and Applied Sciences, University of Pennsylvania, Philadelphia, PA, USA; The Wistar Institute, Philadelphia, PA, USA; Department of Pathology, Perelman School of Medicine, University of Pennsylvania, Philadelphia, PA 19146, USA; Genetics and Epigenetics, Cell and Molecular Biology Graduate Group, Perelman School of Medicine, University of Pennsylvania, Philadelphia, PA, USA; Department of Genetics, Perelman School of Medicine, University of Pennsylvania, Philadelphia, PA, USA

## Abstract

We report a cellular barcoding strategy, SpaceBar, that enables simultaneous clone tracing and spatial transcriptomics profiling. Our approach uses a library of 96 synthetic barcode sequences that can be robustly detected by imaging based spatial transcriptomics (seqFISH), delivered such that each cell is labeled with a combination of barcodes. We used these barcodes to label melanoma cells in a tumor xenograft model and profiled both clone identity and spatial gene expression *in situ*. We developed a gene scoring metric that quantifies how strongly gene expression is driven by intrinsic cellular cues or extrinsic environmental signals. Our framework distinguishes between clonal dynamics and environmentally-driven transcriptional regulation in complex tissue contexts.

## MAIN TEXT

The location of cells in tissues is often tightly linked with their function^1,2^. This realization has led to the development of new technologies for spatial transcriptomic profiling. Additionally, some cellular traits are intrinsically determined, and there is a growing appreciation that a cell’s clonal or lineage identity informs its function^3,4^. However, measuring the relative contributions of cell-intrinsic and environmental factors to a cell’s transcriptomic state has been difficult due to a lack of methods that provide both clonal and spatial information.

The two main types of spatial transcriptomic technologies are sequencing-based and imaging-based. Sequencing-based strategies typically encode spatial information through gridding or beads^5,6^. Traditional clone-tracing technologies, where a cell and its progeny are labelled by integrating a random barcode sequence into the cell of origin and then identified via sequencing, are commonly used with sequencing-based spatial transcriptomics. While these approaches have been informative for constructing broad associations between clone identity, spatial location, and gene expression^7,8^, they are limited by spatial resolution and barcode detection, making it difficult to confidently assign expression patterns to clones at the single-cell level. Imaging-based methods generally offer superior spatial resolution and single-molecule accuracy. However, these methods use pre-determined probe panels, making it challenging to detect the random barcode sequences used in clonal analyses. One approach, Rewind, addresses this challenge by first identifying barcode sequences of interest through sequencing and then detecting them *in situ* using targeted probes^9^, but is not designed for more exploratory work. Alternative approaches, such as MEMOIR^10^, use CRISPR editing of arrays of probe targets to trace lineages and cells, but their complexity and low diversity prevent broader application.

To address these challenges, we developed a system to identify cells of a common origin (clones) together with the transcriptome of cells via imaging-based spatial transcriptomics, which we call SpaceBar. We designed a set of 96 barcode sequences that can be uniquely discriminated with imaging-based transcriptomic technologies such as seqFISH. These barcodes were designed by generating random 1000 base-pair sequences with balanced GC content and minimal repeats or secondary structure. We then constructed a pooled lentiviral expression library of these sequences by inserting these sequences into the 3’ UTR of GFP in a lentiviral backbone, which allowed us to stably introduce these sequences into cells and detect the sequences as RNA transcripts (Figure 1A).

**Figure 1.**
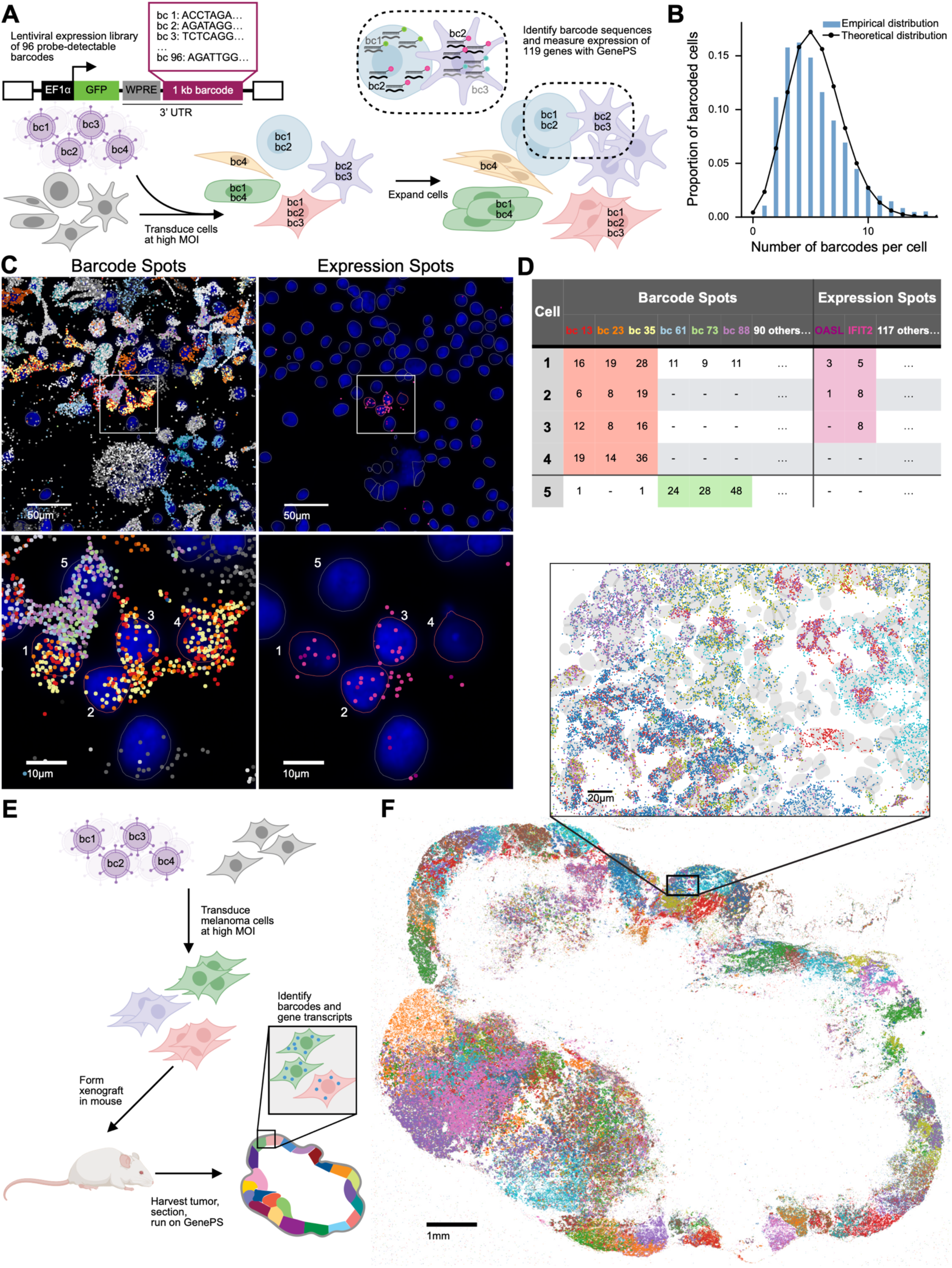
SpaceBar identifies clones *in vitro* and *in vivo*. **A.** Schematic representation of SpaceBar with melanoma cells *in vitro*. To label cells and their progeny, we constructed a pooled lentiviral expression library of 96 1-kb barcode sequences in the 3’ UTR of GFP. We transduced melanoma cells with this library at high MOI such that cells are labeled with a combination of barcode sequences and plated them directly onto a slide. We then expanded the labeled cells and then identified the barcode sequences and the expression of a 120 gene panel using Spatial Genomics’ GenePS system. **B.** The distribution of the number of barcodes detected per cell. The black line depicts a theoretical expectation for the barcode distribution as a Poisson distribution with λ=5.24. **C.** An example of barcode and expression detection of melanoma cells *in vitro* in one field of view. DAPI-stained nuclei are depicted in blue. Nuclear expansions used for spot assignment to cells are depicted with lines. Colored dots indicate the RNA transcript detection. In the left panels, barcode transcripts are depicted such that each color represents a distinct barcode. In the right panels, *IFIT2* and OASL transcripts are depicted in pink. Bottom panels are zoomed-in versions of the top panels showing that cells 1-4 belong to the same clone and also express *IFIT2* and *OASL*. **D.** A cell by gene table of the data shown in C. Each row represents a cell depicted in the image and each column indicates the number of transcript spots assigned to each cell depicted. On the left of the table counts of relevant barcodes per cell, on the right counts for *OASL* and *IFIT2*. **E.** Schematic representation of using SpaceBar with melanoma cells in a xenograft model. We transduced melanoma cells with our lentiviral pool and high MOI. We then expanded these cells and then performed subcutaneous injections to form xenografts in the mouse. After 36 days, tumors were harvested, sectioned, and subsequently run on a GenePS machine for barcode and gene expression detection. **F.** This panel depicts barcode transcripts detected in a section of a tumor from the melanoma xenografts. Each dot depicts a barcode transcript, colored by the barcode identity. Inset shows a subset of the section.

A critical parameter for clone tracing techniques is barcode diversity, which determines the number of individual clones that can be discriminated. We thus transduced cells at a sufficiently high multiplicity of infection (MOI) to ensure cells received unique combinations of the 96 barcodes (Figure 1A). Such a combinatorial scheme could, in principle, scale to uniquely label hundreds of thousands to millions of cells, enabling tissue- or tumor-wide clone analysis through tuning the MOI. For example, for cells that receive at least 3 barcode sequences, there are >140,000 unique combinations of sequences. For applications requiring more diversity, we can achieve over 3,000,000 unique combinations by considering cells with at least 4 barcode sequences. In contrast, for analysis of local clone structure, such as determining whether two nearby cells are from the same lineage, low MOIs using one or two barcodes may be sufficient.

To test our ability to identify cell clones *in vitro*, we transduced 375,000 melanoma cells at an MOI of 5.24, and grew them directly on a slide for spatial transcriptomic analysis. Using Spatial Genomics’ GenePS technology, we used a 216-gene panel to simultaneously detect our library of 96 barcodes and an additional 120 genes of interest, chosen for their relevance to melanoma biology. We profiled approximately 46,000 cells, where 92% had at least one barcode, 64% had three or more, and 47% had at least four (Figure 1B). Of the 120 genes, we removed 7 genes that showed poor detection, resulting in the use of 113 genes for further analysis (see Methods).

We next sought to evaluate the accuracy of our clone identification. A primary challenge with our combinatorial strategy is that two cells may receive the same barcode combination by chance and thus would represent “false clones”. To estimate the rate of false clones, we leveraged spatial information. Given that our cells have limited mobility, we reasoned that neighboring cells with the same barcode combination were likely siblings, whereas distant cells with the same combination of barcodes could represent “false clones” due to independent acquisition of the same barcodes (see Methods for more details). We calculated the fraction of distant cells that shared identical barcodes to estimate the rate of “false clones”. The rate was high for cells with few barcodes—77% for cells with only two barcodes. However, when we restricted our analysis to the 16% of cells with 3 barcodes, the rate dropped to 12% false clones. This rate further dropped to 0.3% when we restricted our analysis to the 47% of cells with at least 4 barcodes (Figure 1D), matching theoretical expectations (Methods, Figure S1B). Thus, our method can uniquely label a high percentage of cells with confidence, and users can tune this degree of confidence by modulating the MOI, number of barcodes, and number of cells included in each experiment (Table S1).

Besides falsely identifying unrelated cells as belonging to the same clone (termed here “false clones”), another type of error occurs when related cells are identified as belonging to different clones (“mis-assignment”). Mis-assignment typically occurs when the barcode is very lowly expressed, not detected, or mistakenly attributed to the wrong cell. An example of mistaken attribution of barcode transcripts is shown in Figure 1C. Cell 1 has 6 barcode transcripts detected and is initially assigned to a clone by itself. However, when examined in its context, it is clear that it contains barcode transcripts from both itself (depicted in yellow, orange, and red) and a neighboring cell 5 (depicted in green, blue, and purple), likely due to overlap in growth. As such, Cell 1 is “mis-assigned” to a clone by itself instead of being appropriately assigned to a clone along with cells 2, 3, and 4, which share the same correct transcripts (yellow, orange, and red). In order to estimate how often this mis-assignment problem occurs, we reasoned that mis-assigned cells would be more likely to share barcodes with neighboring cells not initially assigned to the same clone. Thus, we calculated the fraction of cells that had a high degree of barcode overlap with their non-clone neighbors to estimate the rate of clone mis-assignment. Considering only cells with 4 or more barcodes, 14% of them had at least one neighboring cell that shared at least 4 of the same barcodes (Figure S1). This large overlap suggests that many cells belong to the same clone but are not recognized as such when we use simple presence/absence information to assign clones.

To address this mis-assignment issue, we developed a clustering approach that considers both barcode presence and relative abundance to assign cells to clones. Instead of requiring exact matches, we cluster cells into clones when the relative abundance of barcode sequences are sufficiently similar. This approach improved our clonal assignments to an estimated mis-assignment rate of only 9% (Figure S1). This procedure correctly assigns cell 1 to a clone with cells 2, 3, and 4 in the example shown in Figure 1.

Next, we sought to determine if clonal relationships (as opposed to randomness or external factors) drive gene expression patterns *in vitro*. We first analyzed overall gene expression patterns of the 113 genes detected in our panel and found that cells of the same clone had gene expression profiles that were more similar to each other than random cells or neighboring cells of different clones (p<0.001 as quantified by T-test, see Figure S2). We next wondered if particular genes showed clonal patterns. We first looked at the expression of *OASL* and *IFIT2*, markers of a melanoma cell state that is primed for resistance to targeted therapy and previously shown to have some degree of clonal expression^3,11^). Figure 1C depicts an example of clonal expression of *OASL* and *IFIT2*, where three of the four cells in the clone express one or both of these genes, corroborating the pattern of clonal yet transient expression^3,9,11,12^. We further quantified this pattern, finding that the expression of these pre-resistance marker genes is more likely to be shared between sisters of the same clone than random or neighboring cells (p < 0.001, Fisher’s exact test). This recapitulation of known clonal expression patterns provided strong biological corroboration for our method.

We next tested our method’s ability to detect clonal expansions in tissues. We transduced melanoma cells with our lentiviral barcode pool and generated xenografts in mice (Figure 1E). After 36 days, we harvested tumors and prepared sections for spatial transcriptomics. Using the computational methods we verified on our in vitro sample, we identified clonal populations of cells, demonstrating that our spatial barcodes are suitable for *in vivo* applications (Figure 1F, Figure S3). Here, we focus our analysis on one tissue section containing ∼67,000 cells, with ∼38,000 containing at least one barcode, and ∼16,000 containing three or more barcodes (Figure S4). Using our barcode clustering algorithm, we detected ∼1,400 clones, with an estimated false clone rate of ∼1%.

A major motivation for this method was to quantify how strongly gene expression is determined by spatial context versus intrinsic factors—that is, how much is determined by where the cell is vs. which cells it is related to. Genes with clone-driven expression should have patterns that correspond to clonal relationships (illustrated in Figure 2A), whereas genes with spatially determined expression should have patterns that correspond to the tumor’s spatial structure. Our analyzed tumor section contains a putative vasculature region in its interior (Figure S5) and an exterior characterized by *COL1A1* expression. If this spatial structure drove the expression of a particular gene, we would expect its expression to vary with a cell’s distance from the tumor edge (illustrated in Figure 2B). Alternatively, a gene’s expression could be neither clonally nor spatially determined (Figure 2C).

**Figure 2.**
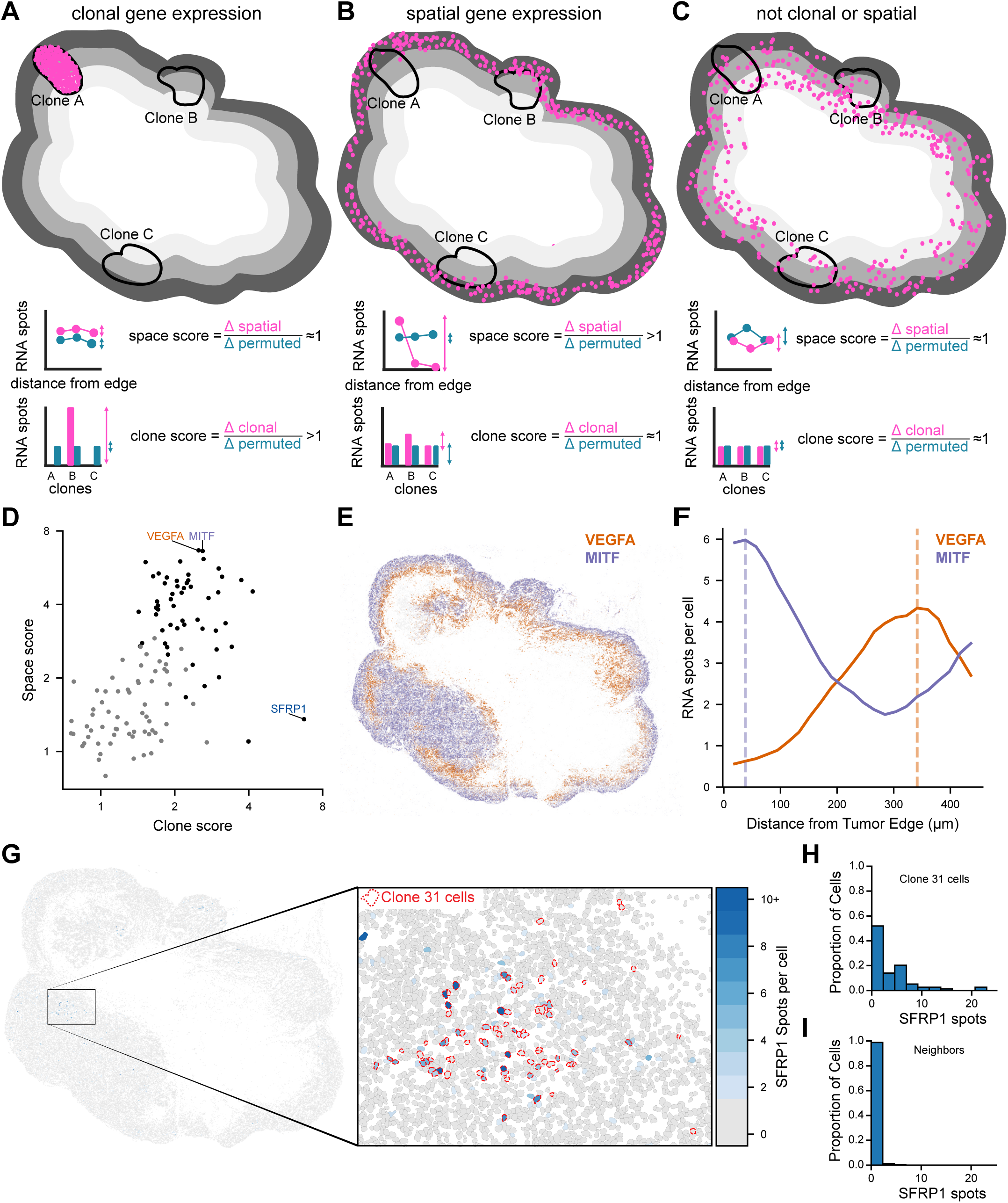
SpaceBar uncovers clonal and spatial gene expression patterns. **A.** Schematic representation of clonal gene expression patterns. The tumor section is divided into groups, with each group representing a clone (depicted by the circles drawn on the tumor). A gene with clonal expression patterns will have higher expression in a particular clone than others, depicted by the pink expression dots in one clone but not the others. This pattern would result in a high clone score, where the difference between the clone with the maximum expression and the clone with the minimum expression is much larger than this difference when clonal identity is permuted. **B.** Schematic representation of spatial gene expression patterns. The tumor section is divided into spatially segmented rings based on the distance from the exterior of the tumor (depicted by gray rings on the tumor). A gene with expression patterns that reflects the interior-exterior spatial structure of this tumor will have varying expression across the spatial segments. This pattern would result in a high space score, where the difference between the spatial segment with the maximum expression and the segment with the minimum expression is much larger than this difference when the cells’ spatial location is permuted. **C.** Schematic representation of gene expression patterns that are not clonal nor spatial. Such a gene would have similar levels of difference between the maximum and minimum expression when the tumor is grouped by clone, spatial distance from the exterior, or when this information is permuted. **D.** A scatterplot of clone and space scores for all 113 genes detected in our tissue section. Each gene is represented by a single point and black points reflect genes with significant clone or space scores, as measured by a clone or space score of at least 2 and a Bonferroni-adjusted empirical p-value < 0.01. Genes with examples depicted in other panels (*VEGFA*, *MITF*, and *SFRP1*) are labeled. **E.** Expression of *MITF* and *VEGFA* in the tumor section. Each dot indicates a detected MITF or *VEGFA* spot, in purple and orange, respectively. **F.** Average expression of *MITF* and *VEGFA* per cell across spatial segments in the tumor section by distance from the exterior of the tumor. *VEGFA* expression is depicted in orange. *MITF* expression is depicted in purple. The vertical dashed lines represent distance from the tumor’s edge with the maximum expression of each gene. **G.** Expression of *SFRP1* in the tumor section. Each cell is colored by the number of *SFRP1* spots detected in the cell. Colorbar is shown in panel H. Cells with at least two *SFRP1* spots are depicted in blue. Cells with zero or one *SFRP1* spots are depicted in gray. The magnification shows the neighborhood of Clone 31, which had the highest mean expression of *SFRP1* of all clones in the tumor section. Clone 31 cells are outlined with a red dashed line. Cells with at least two *SFRP1* spots are depicted in blue (according to the colorbar on the right). Cells with zero or one *SFRP1* spot are depicted in gray. **H.** Distribution of the expression of *SFRP1* across all cells in Clone 31. **I.** Distribution of the expression of *SFRP1* across all cells that are in the neighborhood of Clone 31 (and do not belong to Clone 31).

To determine how much clonal identity or spatial location affected the expression of each gene, we developed two scores (depicted in Figure 2A-C). We binned cells by either clonal identity or by distance from the tumor edge (spatial location). To determine how each gene was associated with clones (clone score) or spatial location (space score), we calculated the average gene expression for each bin (clone or spatial location) and then measured the difference between the bins with maximum and minimum expression. We compared this difference to the average difference from randomly generated bins created by scrambling the clonal identities (or spatial locations). Genes received high clone or space scores if their expression varied more in real versus scrambled bins.

We computed both scores for all 113 genes detected in our panel. 46 genes (40%) had high spatial scores (variation exceeding twice that from scrambled spatial assignment and adjusted p < 0.01). In contrast, only 5 genes (4.4%) showed high clone scores. This difference suggests that, at least for tumors derived from a single recently-bottlenecked cell line, spatial structure is a primary driver of gene expression variation.

Several genes had high space scores, suggesting that their expression patterns corresponded to the internal-external spatial structure of the tumor. *MITF* and *VEGFA* were the two genes with the highest space scores (Figure 2D). *MITF* showed the highest expression levels towards the edge of the tumor, with a peak expression 37 μm from the tumor edge (Figure 2E-F). Conversely, *VEGFA* expression was highest in the interior of the tumor, with a peak expression 334 μm from the tumor edge, showing 6-fold expression in the interior region near the vasculature than the exterior. This pattern is consistent with prior work demonstrating that *VEGFA* promotes the recruitment of vasculature to tumors^13^.

*SFRP1* had the highest clone score, suggesting its expression is clonally determined (Figure 2D). Clone 31 had the highest average *SFRP1* expression, with 3.7 spots per cell, exceeding the tumor-wide average of less than 1 spot per cell (Figure 2E). 58% (46/79) of cells in Clone 31 had non-zero expression of *SFRP1*, exceeding 1.5% (1014/67202) across the tumor or 3.3% (81/2410) of Clone 31’s neighboring cells (Figure 2G-I). Notably, cells that belong to this clone are interspersed with other cells that do not belong to this clone (Figure 2G). Therefore, this clonal expression pattern would be difficult to distinguish from a spatially-driven pattern without sub-cellular resolution (Figure S6).

Our clone score can best detect gene expression patterns with a high degree of heritability. This discriminatory capacity is because the clone sizes in the tumor are large (on average 70 cells per clone), so genes need to be expressed in enough cells to be detected from the clone’s mean expression. This large clone size makes it difficult for our clone score to detect expression patterns that are very transiently heritable, such as the cell state primed for drug resistance identified in the *in vitro* analysis above (Figure 1B,C,D). *In vivo*, these drug resistance priming genes (*IFIT2* and *OASL*) have previously been found in groups neighboring cells in resistance cells post-therapy ^12^, consistent either spatially or clonally-driven gene expression. With our approach, we are able to discern that these groups of cells belong to the same clone, suggesting that the expression of these genes is indeed clonally determined and transiently heritable *in vivo* (Figure S7).

We have developed a cellular barcoding system, SpaceBar, that combines combinatorial labeling using 96 designed barcode sequences with spatial transcriptomics to simultaneously trace clonal relationships and gene expression patterns in tissues. Using this method, we demonstrated that gene expression in tumors is predominantly shaped by spatial organization, with some genes showing strong clonal inheritance patterns. Because our approach relies on commonly used mammalian lentiviral transduction, it can be applied to study a variety of questions in many systems. This technology will be particularly valuable for addressing fundamental questions about how cell-intrinsic and environmental factors influence the function and structure of tissues in cancer, development, regeneration, and beyond.

## Methods

### Cell lines and culture

WM989 melanoma cells (obtained from the laboratory of Meenhard Herlyn) were authenticated via DNA short tandem repeat (STR) microsatellite fingerprinting at the Wistar Institute. Cells were maintained in TU2% media consisting of 78% MCDB (Sigma #M7403), 20% Leibovitz’s L-15 media (Life Technologies Inc., 11415064), 2% FBS, 1.68 mM CaCl2, and supplemented with 50 U/mL penicillin and 50 μg/mL streptomycin (Invitrogen 15140122). To minimize genetic heterogeneity, we utilized WM989-A6-G3 cells, which are single-cell-derived subclones of WM989. Cell passaging was performed using 0.05% trypsin-EDTA (Invitrogen #25300054). All cell lines tested negative for mycoplasma contamination.

### Barcode lentivirus library generation

To design 96 barcode sequences that were uniquely detectable with probes for spatial genomics, we generated 1000 1 kb long sequences with a WSNN pattern to maintain GC content at roughly 50% and avoid long repeats. We then uploaded these sequences to IDT’s complexity score calculator which characterizes sequences in terms of their repeat structure, secondary structure, and suitability for synthesis. We selected the 96 sequences with the lowest complexity scores (all with a score of 1.0 or lower) for synthesis and confirmed their suitability for detection with probes on Spatial Genomics’ GenePS system. We then ordered these 96 sequences as eBlocks from IDT with additional flanking sequences homologous to the backbone for HiFi assembly.

5’ flanking sequence: aagcagtggtatcaacgcagagtacatggg

3’ flanking sequence: gttttagagctagaaatagcaagttaaaataaggc

We constructed a backbone plasmid (pRV27) derived from the CROPseq-Guide-Puro plasmid (Addgene #86708). The puromycin gene was replaced with a MYC-NLS-flanked EGFP derived from pLV[Exp]-EFS>EGFP* (VectorBuilder ID VB211018-1219pmt). Plasmids were linearized via HiFi Phusion PCR (NEB #E0553S) with the following primer sequences: CROPseq-Guide-Puro: ACGCGTTAAGTCGACAATCAACCTCT and CATGGTGGCCGTACGTCACG pLV[Exp]-EFS>EGFP*: TGACGTACGGCCACCATGGTGAGCAAGGG and GTCGACTTAACGCGTTTAATCCAATTTGACGCGCTTTGCAG. These linearized products were joined via In-Fusion cloning (Takara #638948), selected with LB + Carbenicillin, and isolated via miniprep (Qiagen #27104). A TSO site was added to this resulting plasmid (unnecessary for the current application), again via linearization with HiFi Phusion PCR (NEB #E0553S), now with these primers: GTATCAACGCAGAGTACATGGGGACCCAGAGAGGGCCTAT and ACTCTGCGTTGATACCACTGCTTCCCTCGGGGTTGGGA. The linearized product was subsequently joined with In-Fusion cloning (Takara #638948), selected with LB + Carbenicillin, and isolated via miniprep (Qiagen #27104).

To insert barcode sequences into the backbone plasmid, we first performed a restriction digest of pRV27 with Esp3I (NEB #R0734L) and PpuMI (NEB #R0506L). We then used HiFi DNA assembly protocol (NEB #E2621L) to assemble barcode inserts into the digested backbone, with 25.23 ng of barcode insert to 100 ng of digested backbone (1:3 ratio) in a total volume of 20 µL. Assembled product was then transformed into NEB Stable competent cells (NEB #5458) via heat shock and plated on the LB + Carbenicillin plates. After overnight growth at 37 °C, single colonies from each barcode transformant were picked and grown overnight in 3 mL LB at 30 °C, shaking at 200 rpm. 500 µL of saturated culture was mixed with 50% glycerol and subsequently frozen at -80 °C for future construction of the barcode library. Plasmid DNA was extracted from the remainder of the culture using Qiagen Miniprep kit (Qiagen #27104) and whole plasmid sequencing was conducted with Plasmidsaurus for verification.

To construct the plasmid pool, we streaked out all 96 sequence-verified spatial barcode plasmids from glycerol stocks onto LB + Carbenicillin plates. After growth at 37 °C overnight, we picked a single colony from each barcode and grew them at 30 °C in 5 mL of LB + Carbenicillin media for 24 h. After 24 h of growth, we pooled all of the cultures together and performed 4 maxipreps (Qiagen Endofree Maxiprep #12362) on the plasmid pool. We then proceeded to use the plasmid library for lentiviral packaging.

### Lentivirus packaging

We cultured Lenti-X 293T cells (Takara #632180) to confluency in DMEM with 10% FBS medium. Once confluent, we split and counted the cells, plating 15 million cells in 15 cm dishes with the same media. The next day we added 120 µL of polyethylenimine (Polysciences, 23966) to 750 µL of Opti-MEM (Thermo Fisher Scientific, 31985062), separately combining 7.5 µg of VSVG and 11.25 µg of pPAX2 and 15 µg of the spatial barcode plasmid library in 750 µL of Opti-MEM. We then incubated both solutions separately at room temperature for 5 min. We then mixed both solutions together by vortexing and incubated the combined plasmid-polyethylenimine solution at room temperature for 15 min. We added 1658 µL of the combined plasmid-polyethylenimine solution dropwise to eight 15-cm dishes. After 4-5 h, we aspirated the medium from the cells, washed the cells with 1x DPBS, and added fresh TU2% medium. We collected the virus laden media 48 h and 96 h post transfection. We then pooled all lentivirus containing media together and filtered it through a 0.45 µm filter. We used Lenti-X concentrator (Takara #631232) according to the manufacturer instructions to concentrate the virus solution 100-fold. In short, we combined 1 volume of Lenti-X concentrator with 3 volumes of clarified supernatant. We mixed by inversion and incubated overnight at 4 °C. We centrifuged the sample at 1,500 g for 45 minutes at 4°C, a white pellet was visible. We immediately aspirated the supernatant, resuspended in DPBS and stored 50 µL aliquots at -80 °C.

### *In vitro* experiments

To transduce WM989 A6-G3 melanoma cells, we thawed lentivirus aliquots on ice and added it to dissociated melanoma cells such that the lentivirus solution comprised 20% of the total volume of 1000 µL. We plated this suspension containing 375,000 cells directly on a Spatial Genomics flow cell coverslip. We allowed the cells to adhere to the slide overnight and replaced the media the next day with fresh TU2%. We waited a total of 48 h post infection to allow the cells to divide one to two times and then we fixed the sample by incubating it in 4% formaldehyde for 10 min at room temperature, we then washed the slide several times in PBS and incubated the slide in 70% ethanol overnight at 4 °C.

### Mouse xenograft experiment

We created barcoded patient derived xenografts by transducing 1.5 million WM989 melanoma cells, waiting 72 hr to allow the cells to divide 2-3 times and then dissociated the cells. We mixed one million of the dissociated barcoded melanoma cells with growth factor reduced Matrigel (Corning) in a 1:1 ratio and injected the mixture into the right flank of five immunodeficient NSG mice. We allowed the tumors to grow for 36 days before sacrificing the mice, resecting the tumors, and freezing the tumors in OCT on dry ice. The tumors were stored in -80 °C until further processing. We cryosectioned the tumors at 8 µm directly onto a Spatial Genomics compatible slide using a Leica CM3050S cryostat. Immediately after sectioning, we fixed the tumor sections by incubating them in 4% formaldehyde for 10 min at room temperature. We then washed with PBS and incubated the slides in 70% ethanol overnight at 4 °C.

### seqFISH panel design and quality control

The panel was designed to contain probes targeting both the barcode library and genes with interesting expression patterns in melanoma. The melanoma portion of the panel was based on previous work from our lab ^11,12^ and was prepared by curating relevant literature with RNA expression data. Probes for each gene were designed by Spatial Genomics for compatibility with the GenePS machine. The final panel targeted the 96 barcodes in our pooled library and 120 genes of interest.

### Sample preparation and analysis

We prepared and ran the samples using the manufacturer’s protocol. We air dried the sample and incubated in a proprietary clearing solution for 10 minutes. We then removed the clearing solution, rinsed in ethanol and air dried. We denatured primary probes at 90 °C for 3 minutes. At this point, we applied the primary probe solution and placed a parafilm on top of the coverslip to make sure the probe solution completely covered the coverslip. We then incubated the sample at 37 °C overnight in a hydration chamber. The next day, we washed the sample with a primary wash buffer after which we assembled the flow chamber. Once the flow chamber was assembled, we stained the nuclei using rinse buffer with DAPI and then washed the sample several times with rinse buffer. We then loaded the sample into the GenePS machine for rounds of secondary hybridization and readout.

### Data processing, segmentation, and spot assignment

Spots for each region of interest were manually thresholded and subsequently decoded using Spatial Genomics’ analysis software. Nuclei were segmented using Spatial Genomics’ analysis software. We then assigned transcripts to nuclei using the custom-built SGObject package with an expansion of 10 pixels around each nucleus. For expanded nuclei that overlap, transcripts were assigned to the nearest nucleus.

### Data quality control and exclusion of poorly detected genes

We assessed the performance of probes targeting genes in our panel by considering the number of transcripts detected in each cell. We excluded all genes from further consideration that were not detected in at least one cell with at least 5 transcripts in both the *in vivo* and *in vitro* samples we primarily consider in the main text. This resulted in the exclusion of 7 poorly detected genes from our panel: *BMP4*, *ITGA8*, *IGFBP2*, *KIT*, *NANOG*, *ROR2*, and *SOX2*. We used the remaining 113 genes (and all 96 barcodes) for further analysis.

### Barcode identity assignment and clustering

For initial barcode assignment based only on transcript abundance, we considered only barcodes that were detected as at least 3 transcripts in a given cell. We then assigned all cells with the same combination of barcodes detected above this threshold to a clone.

To improve our clone assignment, we devised an approach to cluster cells into clones based on the relative abundance of detected barcodes. We first restricted our analysis to cells with at least 10 detected barcode transcripts. We then computed the full pairwise Bray-Curtis dissimilarity matrix based on barcode counts per cell normalized to the total number of barcode spots assigned to that cell. Cells were then clustered together using Agglomerative Clustering with the scikit-learn python package, with clusters merged together based on the average pairwise Bray-Curtis dissimilarity. We chose a threshold value of 0.2 for clone assignments in the *in vitro* experiments and a threshold value of 0.4 for clone assignments *in vivo*, based on manual verification of clustering results.

### Barcode identification error detection

We used two measures to evaluate our clone identification procedure. We first asked how often unrelated cells received the same combination of barcodes by chance (and are “false clones”). To estimate the “false clone” rate, we reasoned that nearby cells with the same combination of barcodes are likely to be descendants from the same initially labeled cells whereas distant cells with the same combination of barcodes are likely to be false clones. We chose a distance of 287 µm as our threshold for “distant cells”. This distance represents the 99th percentile of pairwise distance with cells of the same combination of barcodes for cells with a large number of barcodes (4 or more) (Figure S1). We then calculated the fraction of barcode combinations that had at least two cells greater than this threshold distance apart as our “false clone” rate.

We also computed a theoretical expectation for the false clone rate. This expectation is the expected number of cells expected to share the same combination of barcodes based on the number of cells initially labelled with barcodes. This expectation is given by:

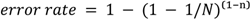

where N = (96 choose k) is the binomial coefficient describing the number of possible barcode combinations for cells that received *k* barcodes from the total barcode library size of 96. The variable *n* denotes the number of initially barcoded cells (clones) in the population. This formula accounts for the probability of unique barcode assignment across the cellular population (Fig. S1B).

We also calculated a “mis-assignment” rate, which reflects cells that are incorrectly assigned to a clone by itself (or with non-clonal neighbors) due to low barcode expression, poor detection, or mis-attributed spots due to cell overlap. To quantify this error, we analyzed the relationship between cells and their spatial neighbors (defined as cells within 30 μm, yielding an average of five neighboring cells). For each cell, we calculated the fraction of non-clonal neighbors that shared barcodes with it. We then iteratively tested different barcode-sharing thresholds to understand how the mis-assignment rate varied with the stringency of our clonal grouping criteria. This analysis helped us evaluate cases where cells might be incorrectly excluded from their true clone or where a single clone might be artificially split into several distinct groups due to imperfect barcode detection.

### Calculating clone and space scores

To calculate the clone score for the *in vivo* data, we first excluded our analysis to clones with at least 3 assigned barcodes and clones with at least 25 cells to prevent very small clones and clones that happened to receive the same barcode combinations from driving the clone score calculation. Then, for each gene, we computed its mean expression in each clone and subsequently calculated the difference between its maximum mean expression across clones and the minimum mean expression across clones (Δ clonal). We then scrambled the clone identity of cells and computed the same difference 12,000 times, taking the average difference as Δ permuted. We then computed the ratio of the gene’s difference across clones to the gene’s difference across permuted clonal identity as the clone score:

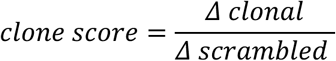

To test whether the clone scores significantly deviated from 1, we computed a Bonferroni-corrected empirical p-value based on the number of instances the difference across scrambled clones met or exceeded the difference from real clone identity (Δ clonal).

To calculate space scores, we binned cells by their distance from the tumor edge. Specifically, we created 25 bins, each ∼18.6 μm in width, from the edge of the tumor. We also included only cells that had at least one barcode detected, to ensure that mouse cells were excluded from the analysis. We then computed an analogous score to the clone score, instead using the spatial bins to group cells instead of the clones. Specifically, we computed the difference between the bin with the highest mean expression and the bin with the lowest mean expression for each gene (Δ spatial). We then scrambled the spatial bins of the cells and computed the same difference 12,000 times, taking the average difference as Δ permuted. We then computed the ratio of the gene’s difference across spatial bins to the gene’s difference across scrambled spatial locations as the space score:

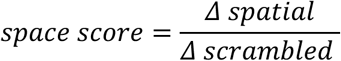

As in the clone score, we computed a Bonferroni-corrected empirical p-value based on the number of times the scrambled spatial bins met or exceeded the difference identified by the real spatial locations to determine significance.

### Verification of spatial expression of *VEGFA* and *MITF*

To validate and quantify the spatial expression of *VEGFA* and *MITF*, the two genes receiving highest spatial scores from our analysis, we calculated the mean number of transcripts per cell across the 25 spatial bins. To smooth out noise from our data, we calculated the average using moving windows of size 3.

### Verification of clonal expression of *SFRP1*

To validate and quantify the clonal expression of *SFRP1*, which had the highest clone score in our analysis, we first examined the clones with the highest mean *SFRP1* expression. We found one clone, Clone 31 that had much higher expression than other clones, with a mean expression of 3.4 transcripts per cell. We then examined the neighborhood of this clone for the expression of barcodes. Upon further examination, we identified some cells that belonged to the same clone but were not clustered together with Clone 31, primarily due to the presence of barcode transcripts from neighboring cells. We performed manual clustering of this cluster to ensure complete assignment of cells appropriately to this clone. This manual clustering was conducted without information about *SFRP1* expression in each cell. We then examined the expression of *SFRP1* from this manually-curated Clone 31, its neighbors, and the entire tumor, finding a significant enrichment of *SFRP1* expression in this tumor.

To understand the importance of sub-cellular resolution to the identification of *SFRP1*’s clonal expression pattern, we also conducted a pseudo-gridding procedure to divide expression spots into 55 μm grids, similar in resolution to Visium’s expression spots. To simulate clone detection with this resolution, we considered all clones with at least 10 cells in this neighborhood and summed the barcode transcripts for all transcripts associated with the clone, treating all of them as a label for the clone. From this analysis, we found that while Clone 31 does indeed have the highest association with *SFRP1* expression (as confirmed by our sub-cellular resolution analysis), eight other clones had positive and significant correlation with *SFRP1* expression, which would have made it difficult to definitively show that *SFRP1*’s expression was indeed clonal and not merely spatially-determined (Fig S6).

## Code and data availability

Code for analysis and figure generation and processed data are available on GitHub: https://github.com/grantkinsler/SpatialBarcodes. Raw data is available upon request to the authors.

## Supporting information

Supplemental Table - seqFISH panel

Supplemental Table - Barcode sequences

## Acknowledgments

We thank Catherine Triandafillou and members of the Raj lab for helpful discussions and feedback on the work. G.K. acknowledges support from NIH Training Grant T32-CA-009140. A.R. acknowledges support from the Samuel Waxman Cancer Research Foundation, the Mark Foundation, and the Melanoma Research Alliance. R.H.B. acknowledges support from NIH Training Grant in Computational Genomics T32HG000046 and NIH Medical Scientist Training Program T32GM007170.

## Declaration of interests

A.R. receives royalties related to Stellaris RNA FISH probes. A.R. serves on the scientific advisory board of Spatial Genomics. All other authors declare no competing interests.

## Supplemental Figures

**Figure S1.**
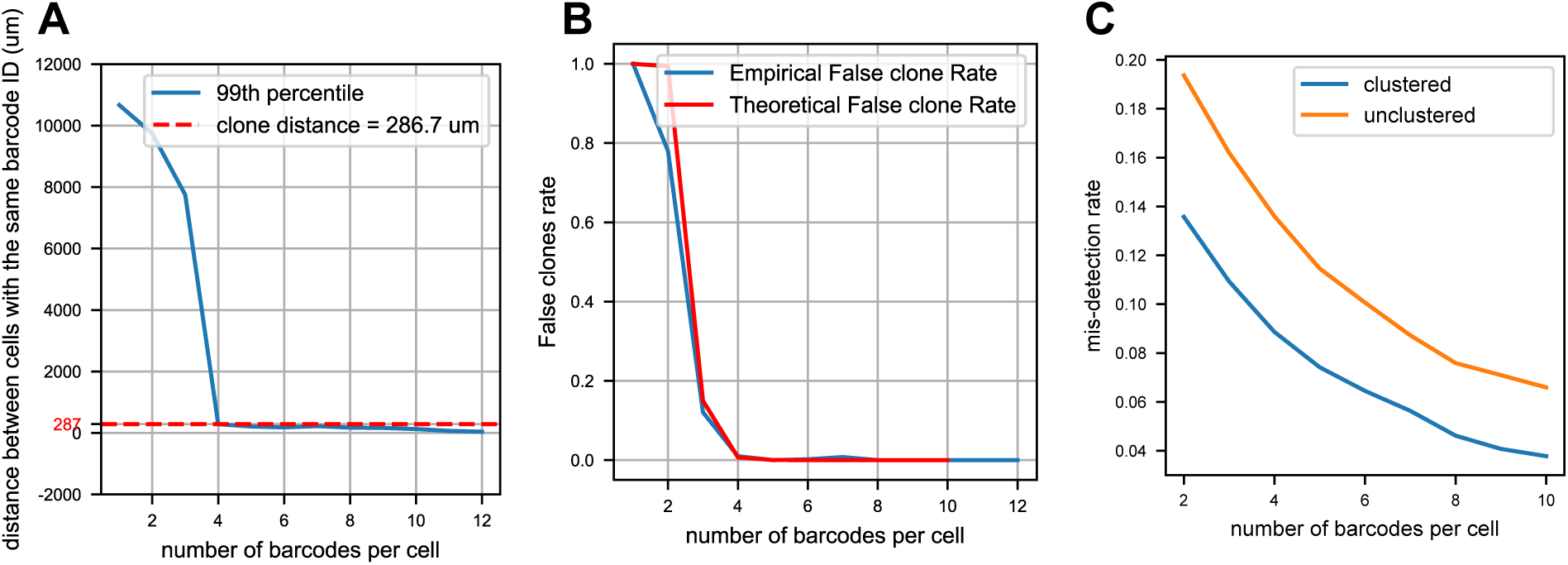
Uniqueness of barcode combinations, clustering reduces barcode mis-detection. A. Top 99th percentile distance between clone cells vs the number of barcodes integrated per cell. We use the value for four barcodes as the distance threshold above which the clone calling would be considered “false clones”. B. “False clone” rate reflected by the frequency of clone with above threshold size. C. Comparing the mis-detection rate (reflected by neighbor barcode overlap) between clustered and unclustered data vs number of barcodes per cell

**Table S1.** False clone detection variation with number of barcodes integrated per cell and number of initial clones transduced.

| Region | Number of clones | False clone rate for cells with corresponding number of barcodes |  |  |
| --- | --- | --- | --- | --- |
|  |  | 2 barcodes | 3 barcodes | 4+ barcodes |
| ROI 2 | 8,898 | 45% | 3.4% | 0.2% |
| All ROIs | 23,188 | 78% | 12% | 0.4% |

**Figure S2.**
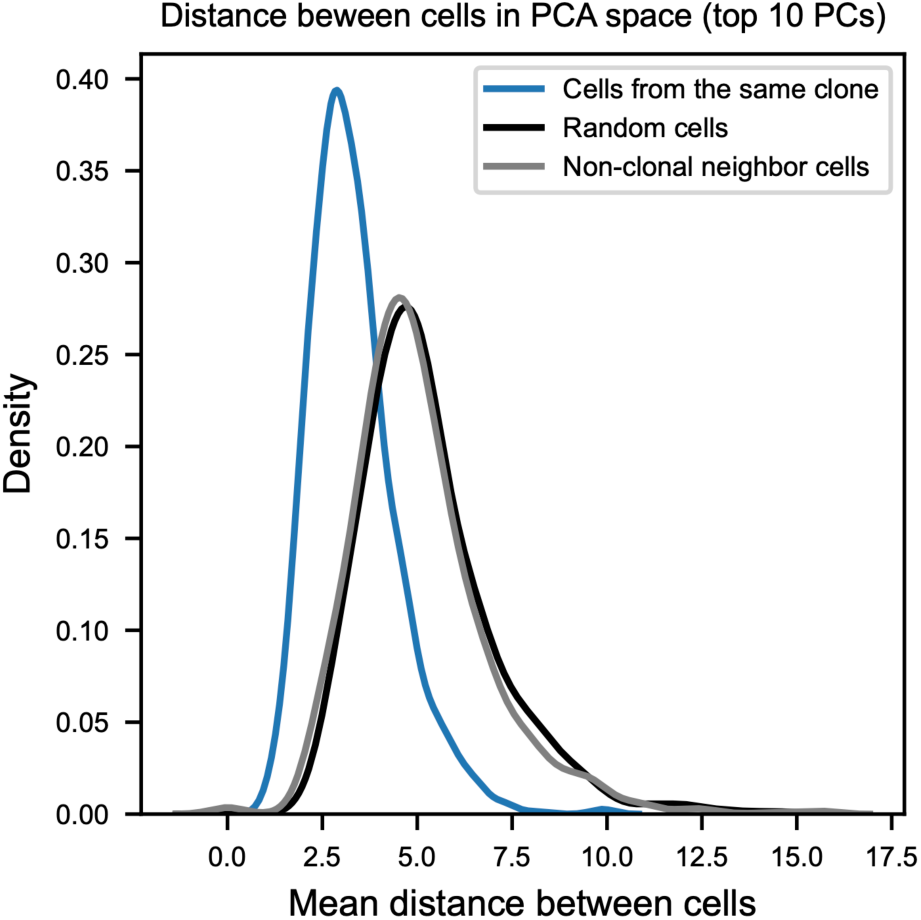
*In vitro,* cells of the same clone have more similar gene expression than random cells. Depicts histograms for the mean pairwise distance in PCA space (of the top 10 PCs) of cells that belong to the same clone. Blue histogram shows the main pairwise distance for actual clones. Gray histogram depicts cells randomly assigned to clones (keeping the number of clones and cells per clone as the real distribution).

**Figure S3.**
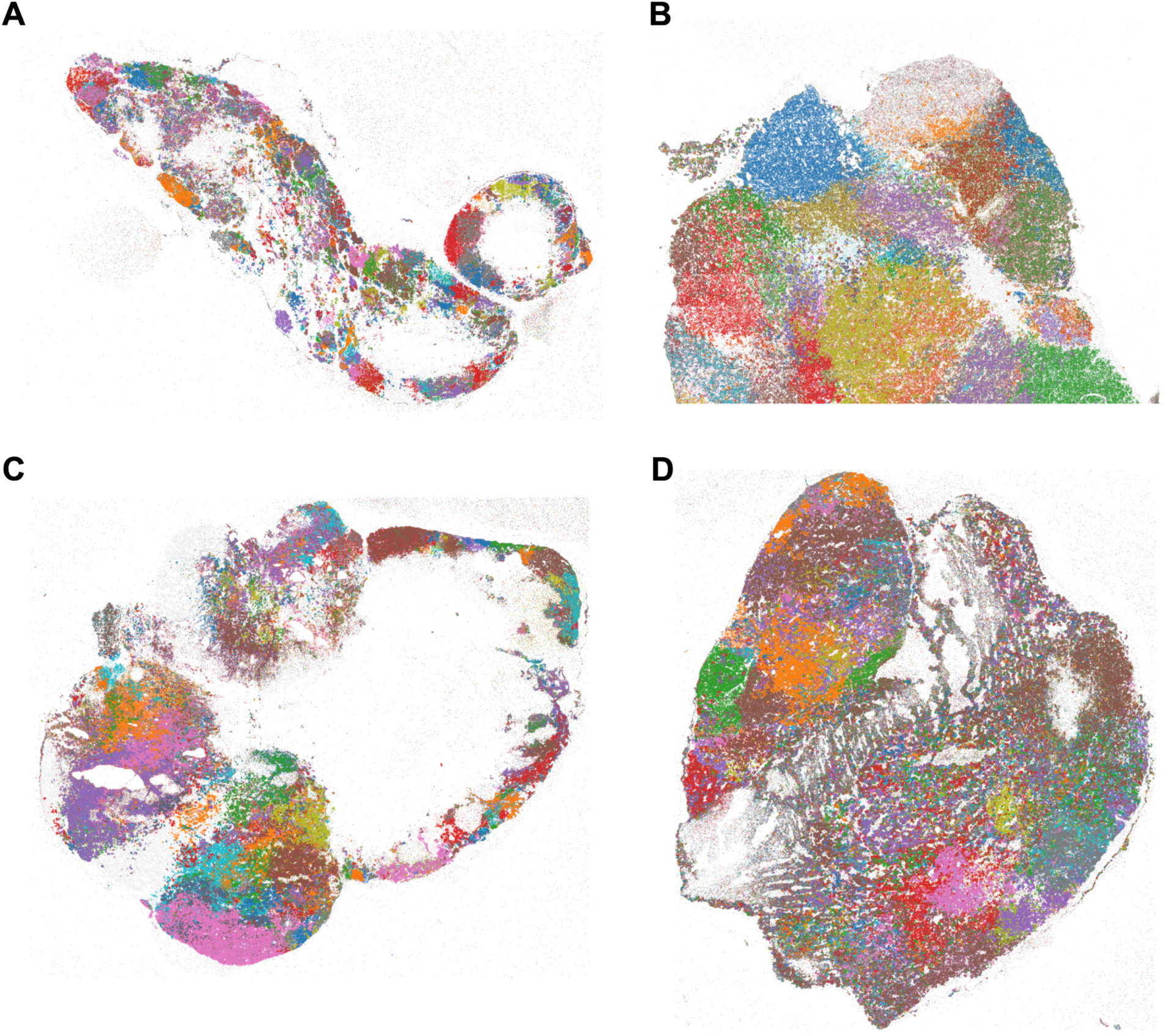
SpaceBar identifies clonal structure in several xenograft tumors. Each subpanel depicts a section of a melanoma xenograft from a different mouse. Each dot depicts a barcode transcript, colored by the barcode identity. Subpanel C is a separate section from the same tumor as the section primarily considered in the main text.

**Figure S4.**
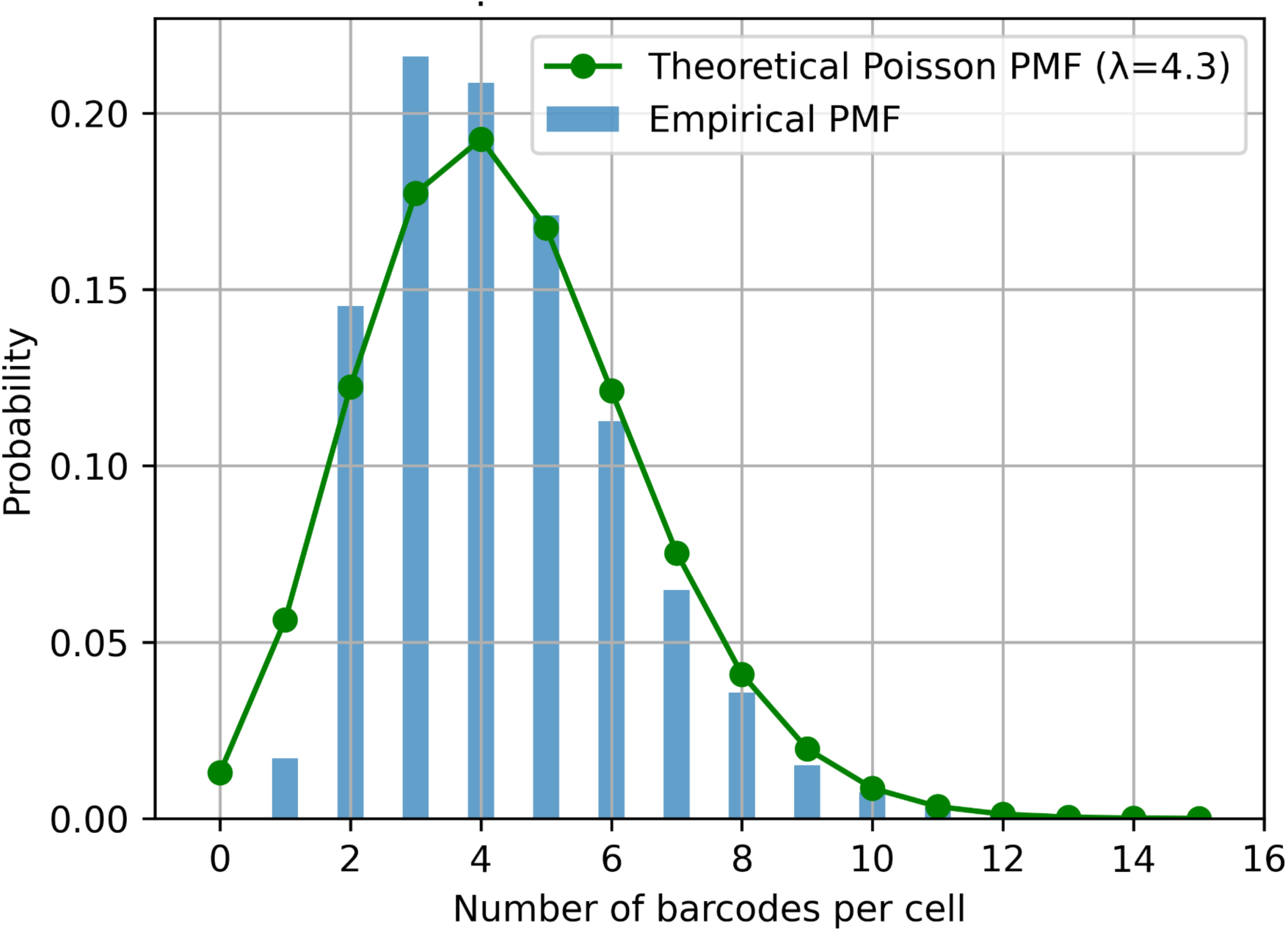
Number of barcodes per cell *in vivo*. This histogram represents the number of barcodes detected per cell in the xenograft section, using a minimum count of 3 spots for a given barcode. Green line represents the theoretical expectation given by a Poisson distribution with λ=4.3, which is the mean for the section.

**Figure S5.**
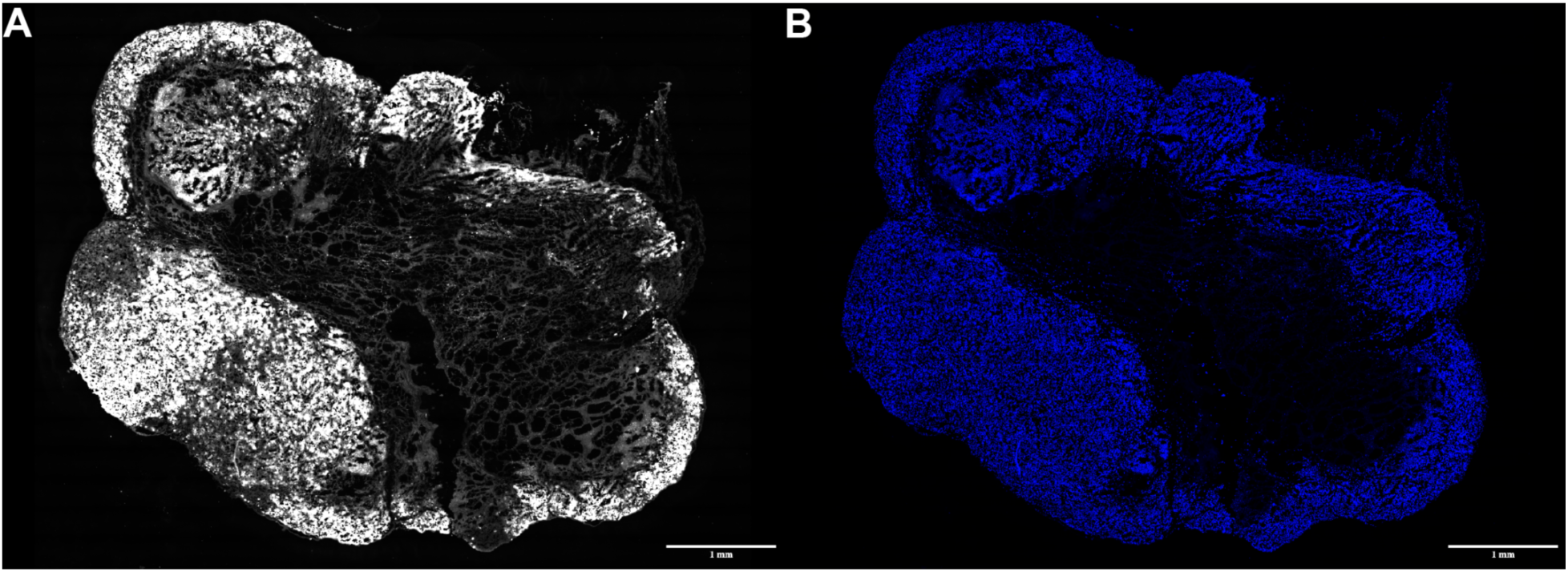
Tumor section images. **A.** GFP channel from the first hybridization round of the focal xenograft section. Barcoded cells are GFP+ and autofluorescence shows the vascularized interior of the tumor. **B.** DAPI channel of the section.

**Figure S6.**
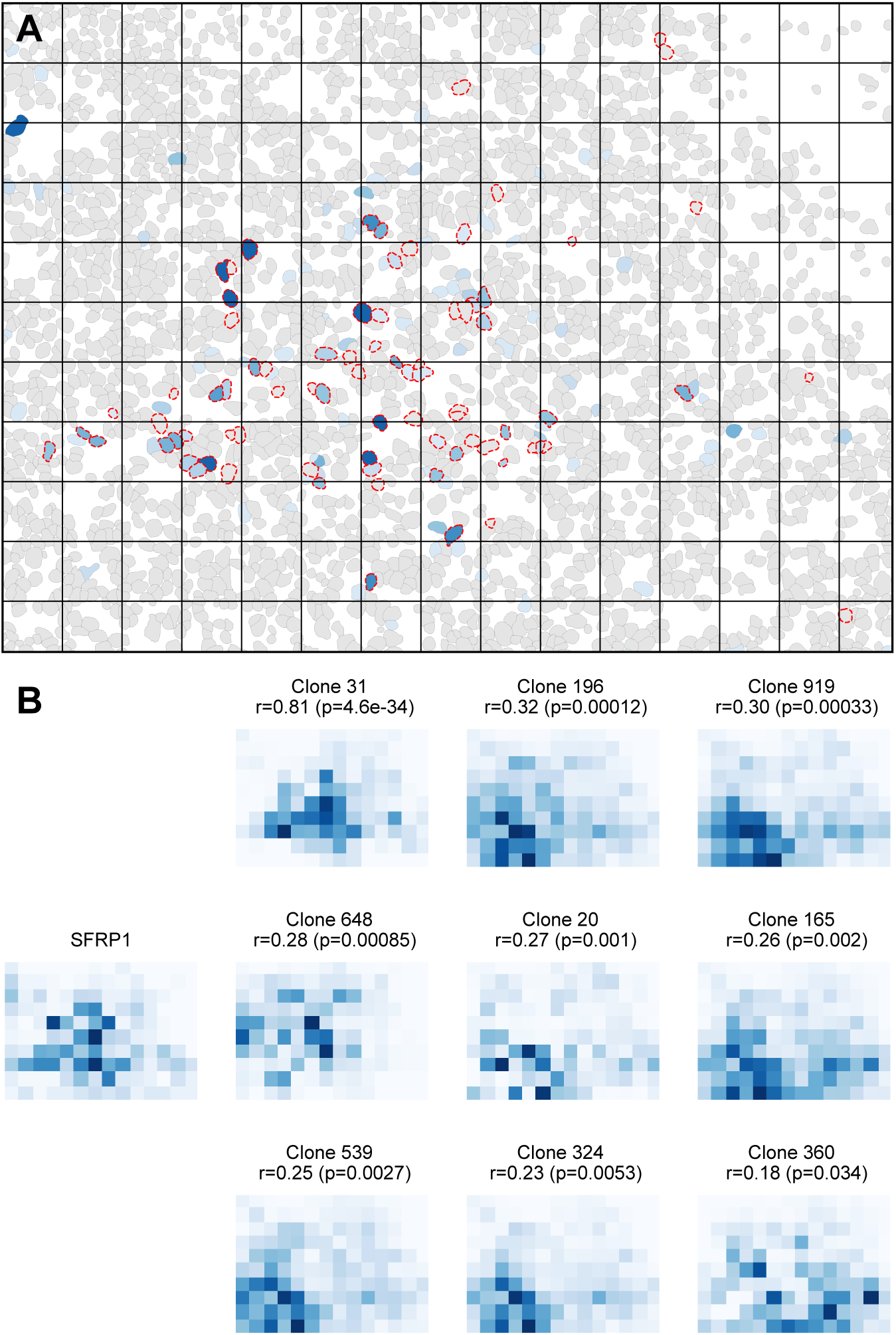
*SFRP1* clonal structure would be difficult to detect without high resolution. **A.** Expression of *SFRP1* in the neighborhood of Clone 31 as depicted in Figure 2, with grids spaced 55um apart. **B.** A series of heatmaps that depict the expression pattern of *SFRP1* as well as the distribution of spots that label distinct clones in this area of the tumor section. Clones depicted as the sum of all transcripts for barcodes assigned to that clone. All nine depicted clones have significant correlations (p < 0.01) with *SFRP1* expression across the grid.

**Figure S7.**
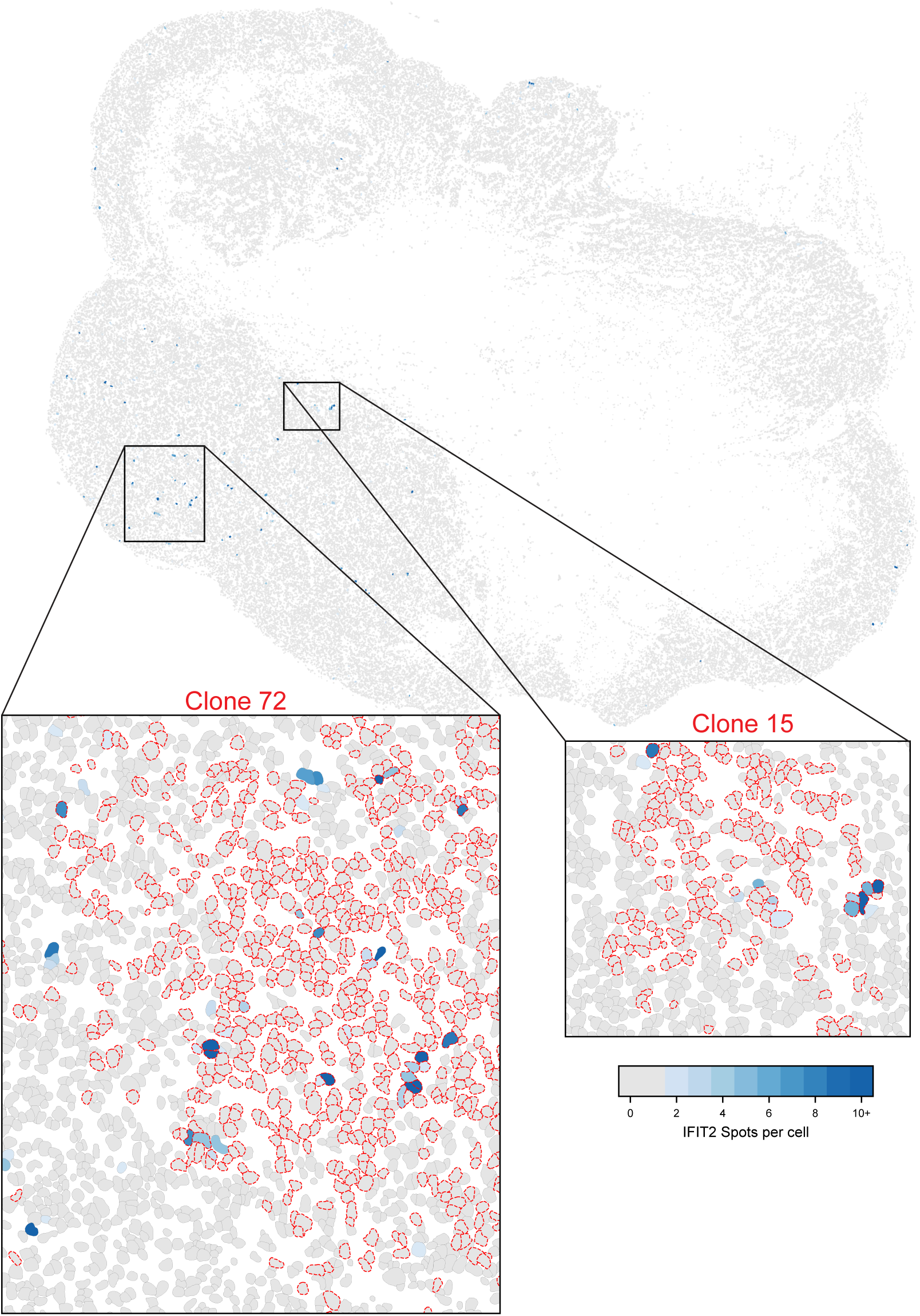
*IFIT2* has high expression in nearby cells of the same clone, suggesting transiently heritable expression. The expression of *IFIT2* is shown in the tumor section. Color indicates the expression level per cell; cells with zero or one spot are depicted in gray. The first inset depicts the expression of *IFIT2* in the neighborhood of Clone 72 (clone 72 cells outlined in red). The second inset depicts the expression of *IFIT2* in the neighborhood of Clone 15 (clone 15 cells outlined in red). *IFIT2* expression is high in groups of 3-5 cells within the same clone, suggesting its expression is transiently heritable.

